# The PTPRT pseudo-phosphatase domain is a denitrase

**DOI:** 10.1101/238980

**Authors:** Yiqing Zhao, Shuliang Zhao, Yujun Hao, Benlian Wang, Peng Zhang, Xiujing Feng, Yicheng Chen, Jing Song, John Mieyal, Sanford Markowitz, Rob M. Ewing, Ronald A. Conlon, Masaru Miyagi, Zhenghe Wang

## Abstract

Protein tyrosine nitration occurs under both physiological and pathological conditions^1^. However, enzymes that remove this protein modification have not yet been identified. Here we report that the pseudo-phosphatase domain of protein tyrosine receptor T (PTPRT) is a denitrase that removes nitro-groups from tyrosine residues in paxillin. PTPRT normally functions as a tumor suppressor and is frequently mutated in a variety of human cancers including colorectal cancer^2,3^. We demonstrate that some of the tumor-derived mutations located in the pseudophosphatase domain impair the denitrase activity. Moreover, PTPRT mutant mice that inactivate the denitrase activity are susceptible to carcinogen-induced colon tumor formation. This study uncovers a novel enzymatic activity that is involved in tumor suppression.

---

The human genome encodes 21 receptor protein tyrosine phosphatases (RPTPs)^4,5^. Twelve of the 21 RPTPs have two catalytic PTP domains in their intracellular parts^3,4^. While the membrane proximal PTP domains (D1) are protein tyrosine phosphatases, it has been thought that the C-terminal PTP domains (D2) lack enzymatic activity^6,7^. Protein tyrosine phosphatase receptor type T (PTPRT) is frequently mutated in human cancers including colorectal cancer^2,3^. We demonstrated that PTPRT knockout (KO) mice are susceptible to colon cancer^8,9^, thereby functioning as a tumor suppressor. Interestingly, one fifth of tumor-derived mutations of PTPRT are located in the D2 pseudo-PTP domain3, consistent with the D2 domain also having tumor suppressor activity.

We hypothesized that the D2 domain of PTPRT may catalyze removal of other types of tyrosine modifications. In addition to phosphorylation, tyrosine residues in proteins can undergo nitration at the 3-carbon position on the phenol ring (Y-NO2) or sulfation of the hydroxyl group10. To test whether PTPRT regulates these modifications, we blotted brain lysates of a PTPRT KO mouse and its WT littermate with antibodies against either nitrotyrosine (N-Y) or sulfotyrosine (S-Y).

As shown in Fig. 1A, nitrotyrosine levels of several proteins were higher in the brain lysate of PTPRT KO mouse compared to the WT littermate, whereas no change in protein tyrosine sulfation was observed (Fig. 1B). Our attempts to identify the modified proteins were unsuccessful, but given that all D2 domains of RPTPs co-exist with the PTP domains (D1), we postulated that the tyrosine phosphatase domain (D1) could share substrates with the D2 domain. Because previously we had identified paxillin and STAT3 as phosphatase substrates of PTPRT^8,11^, we tested if PTPRT regulates nitration of these proteins. As shown in Fig. 1C, nitrotyrosine levels of paxillin, but not STAT3 (data not shown), were increased in brain lysates of PTPRT KO mice compared to the WT littermates. Conversely, compared to GFP control, overexpression of WT PTPRT, but not a PTPRT construct devoid of the D2 domain (Fig. 1D), decreases nitrotyrosine levels of paxillin (Fig. 1E). Together, our data suggest that the D2 domain of PTPRT is responsible for the reduced paxillin tyrosine nitration.

**Figure 1.**
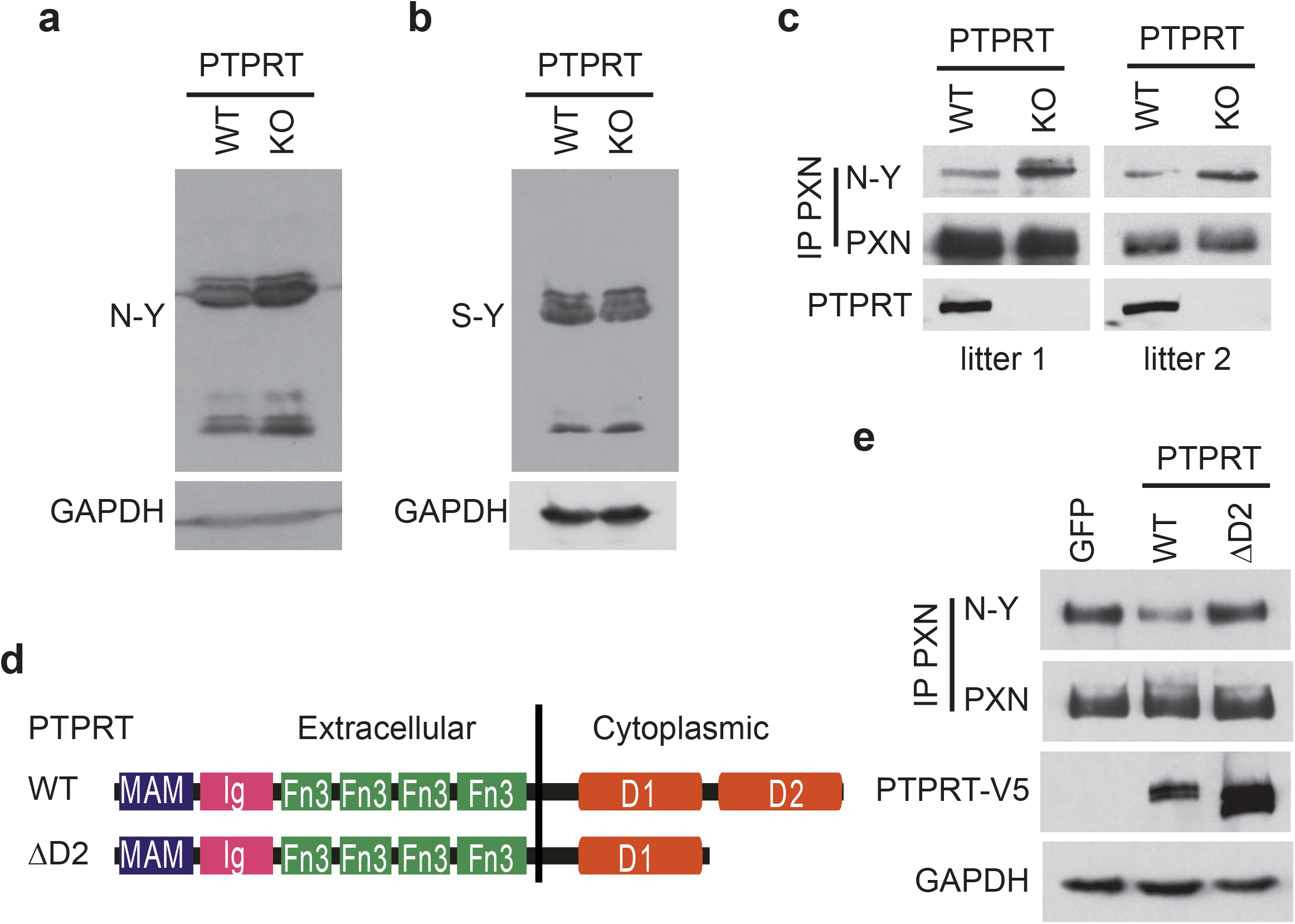
The second catalytic domain (D2) of PTPRT regulates paxillin (PXN) tyrosine nitration (N-Y). (A) & (B) *Levels of tyrosine nitration, but not sulfation, are increased in PTPRT KO mice.* Brain lysates of mice with the indicated genotype was blotted with an antibody against either nitrotyrosine (A) or sulfotyrosine (B). (**C**) *Levels of paxillin tyrosine nitration are increased in PTPRT KO mice.* Brain lysates of littermates of the indicated genotypes were immunoprecipitated with an anti-paxillin (PXN) antibody. The immunocomplexes were blotted with the indicated antibody. Brain lysates were also blotted with an anti-PTPRT antibody. (**D**) *Schematics of PTPRT constructs*: WT: wild-type; ΔD2: PTPRT devoid of the second catalytic domain (D2). (**E**) *The D2 domain of PTPRT regulates paxillin tyrosine nitration.* HCT116 cells were infected with adenoviruses expressing GFP, V5-tagged WT PTPRT, or ΔD2. Cell lysates were immunoprecipitated with an anti-paxillin antibody and blotted with an anti-N-Y antibody. Cell lysates were also blotted with the indicated antibodies.

We further demonstrated that the recombinant D2 domain of PTPRT, purified from *E. coli*, directly decreases nitrotyrosine levels of full-length paxillin immunopreciptated from cell lysates in vitro (Fig. 2 A and B). A D2 C1400A mutant in which the cysteine cognate to the highly conserved catalytic motif of D1 phosphatases is mutated had no activity in this assay. Using a series of paxillin deletion constructs (Fig 2 A, C and D), we mapped two tyrosine residues (Y404 and Y409) in the middle part of paxillin (M-2) as substrates of the PTPRT D2 domain. Mass spectrometry analyses demonstrated *in vitro* denitration of the two tyrosine residues in paxillin by the D2 domain of PTPRT (Fig. 2E and Extended Data Fig. 1 & 2). In contrast, the D1 domain of PTPRT did not affect nitrotyrosine levels of the same substrate (Fig. 2F). Moreover, the purified D2 domain decreases nitrotyrosine levels of the paxillin M-2 fragment in a time-dependent manner (Fig. 2G). Taken together, these data suggest that the PTPRT pseudo-PTP domain (D2) possesses denitrase activity that removes nitro-groups from tyrosine residues in paxillin.

**Figure 2.**
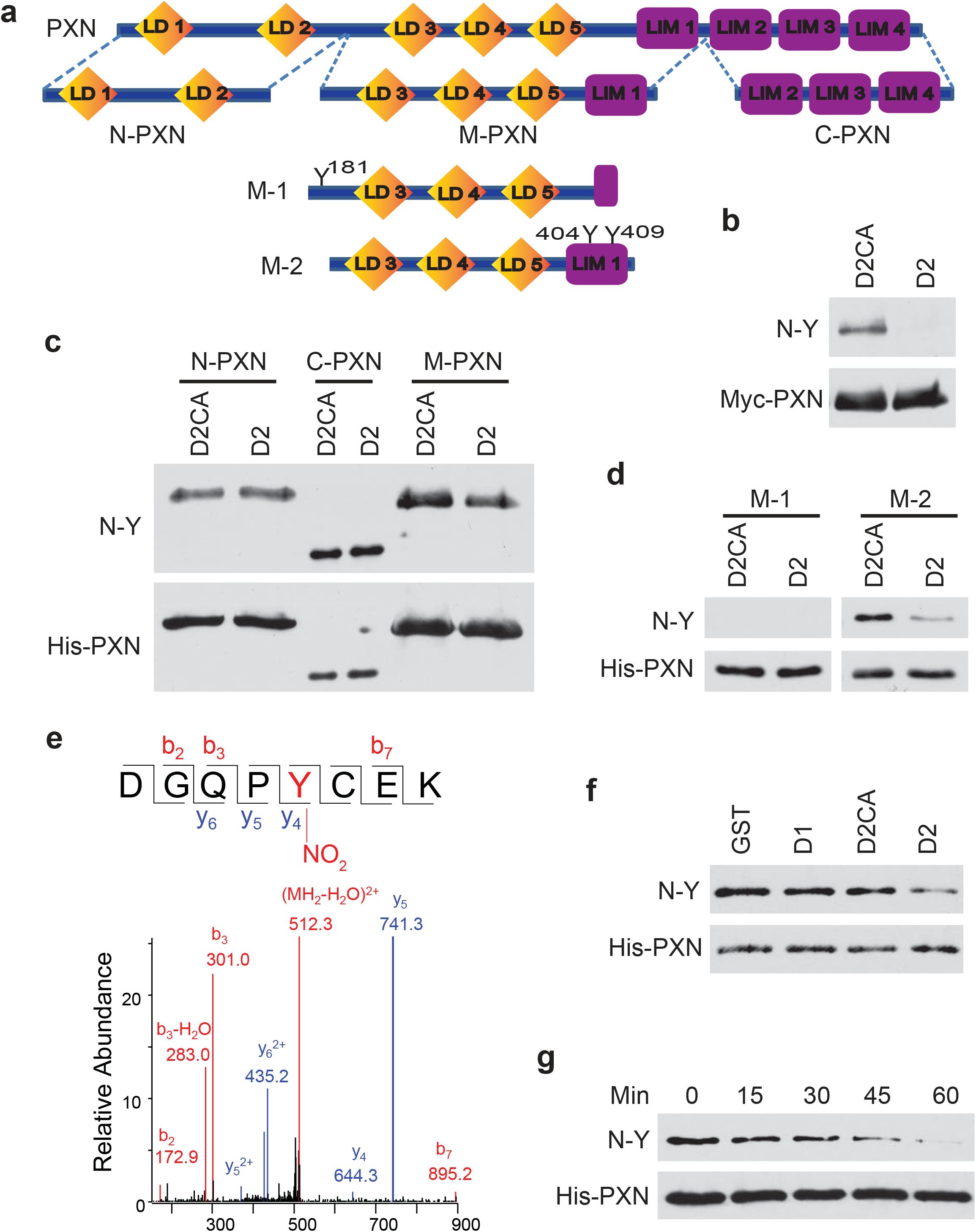
The D2 domain of PTPRT removes paxillin tyrosine nitration *in vitro.* (A) *Schematics of paxillin full-length and deletion constructs.* (**B**) *Recombinant PTPRT D2 domain removes paxillin tyrosine nitration.* HCT116 cells expressing Myc-tagged paxillin were treated with peroxynitrate and paxillin proteins were immunoprecipitated as substrate and mixed with either recombinant WT D2 or D2 C1400A (D2CA) mutant proteins to perform *in vitro* denitrase assay (see supplemental methods section for details). The reaction mixtures were resolved on a SDS-PAGE gel and Western bots were performed with the indicated antibodies. (**C**) & (**D**) *The D2 domain of PTPRT removes tyrosine nitration at residues 404 and 409 in paxillin.* The indicated recombinant paxillin fragments were nitrated by tetranitromethane (TNM) as substrates for the *in vitro* denitrase assays. (**E**) *Validation of paxillin tyrosine denitration by mass spectrometry analyses.* The nitrated paxillin M-2 fragments were treated as shown in (D) and resolved on a SDS-PAGE gel. The paxillin M-2 bands were excised for mass spectrometry analyses. MS/MS spectra of a nitro-Y404 paxillin peptide (DGQPYCEK, residue 400-407) are shown in (**E**). Doubly charged precursor ions from the nitrated (m/z 521.2) and denitrated (m/z 498.7) peptides were subjected to collision-induced dissociation. Cys405 in the peptide was carbamidemethylated. A series of b and y series fragment ions that verify the chemical structures of the peptides were observed. Quantification of nitrated paxillin Y404 and Y409 is shown in supplementary Figures 1 and 2. (**F**) *The PTPRT D1 domain has no denitrase activity.* The indicated PTPRT recombinant protein or GST was mixed with nitrated paxillin M-2 fragment to perform the *in vitro* denitrase assay. (**G**) *Time-dependent denitration of paxillin M-2 by the PTPRT D2 domain.*

To determine the enzyme kinetics of the PTPRT denitrase, we used a synthetic nitrotyrosine-containing paxillin peptide encompassing the nitrated Y404 residue as the substrate. As shown in Fig. 3 A & B, the WT D2 domain has Km of 1.6 mM and kcat of 0.37 S^-1^, whereas the D2 domain with a mutation in the putative catalytic cysteine (C1400A) has no detectable enzymatic activity.

We tested 4 recurrent tumor-derived D2 mutations for denitrase activity. Interesting, all 4 mutant proteins (P1235L, E1299G, G1394R and R1046C) had an increased Km (Fig. 3B), suggesting that these mutations impair binding of substrates to the D2 domain. Of note, all WT and mutant PTPRT D2 proteins were purified to near homogeneity (Fig. 3C).

**Figure 3.**
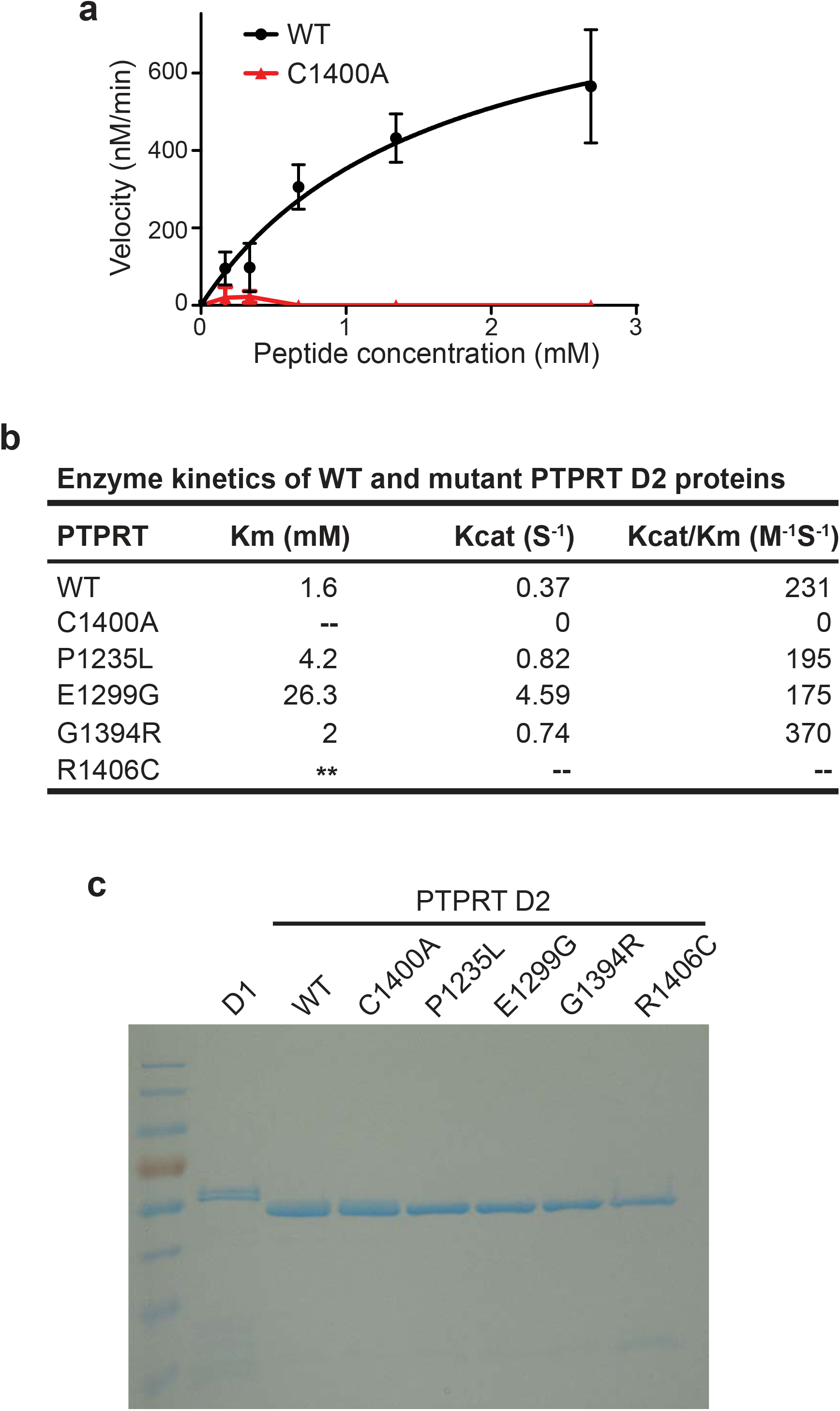
Tumor-derived D2 domain mutations of PTPRT impair the denitrase activity. The indicated WT or mutant D2 recombinant proteins were mixed with various concentrations of a synthesized nitrated Y404 paxillin peptide for enzyme kinetics analyses (see supplementary methods for details). The kinetic curves of D2 WT and C1400A mutant are shown in (**A**) and kinetics parameters of the indicated proteins are shown in (**B**); -- indicates indeterminable; ** indicates too large to determine (some activity was observed at high substrate concentration, but substrate solubility precluded study of concentration-dependence). Purified recombinant D1, WT D2 and mutant D2 proteins are shown in (**C**).

To test if the PTPRT denitrase contributed to tumor suppression, we generated Ptprt^C1400V^ mutant knockin mice in the C57BL/6 mouse strain using CRISPR/Cas9 genome editing (Fig. 4A and Supplementary Fig. 3). We chose to mutate the catalytic cysteine to valine, because it creates a new PmlI restriction cutting site that facilitates genotyping of the mutant mice (Extended Data Fig. 3). Nitrotyrosine levels of paxillin were dramatically up-regulated in Ptprt^C1400V/C1400V^ mice compared to their WT littermates (Fig. 4B). The C1400V mutation did not affect the phosphatase activity, as the levels of pSTAT3, a known phosphatase substrate of PTPRT, did not change in Ptprt^C1400V/C1400V^ mice compared to the WT littermates (Fig. 4C). The mutation also did not affect levels of PTPRT protein (Fig. 4D). Together, these data provide compelling evidence that the denitrase activity of PTPRT decreases paxillin tyrosine nitration *in vivo.*

**Figure 4.**
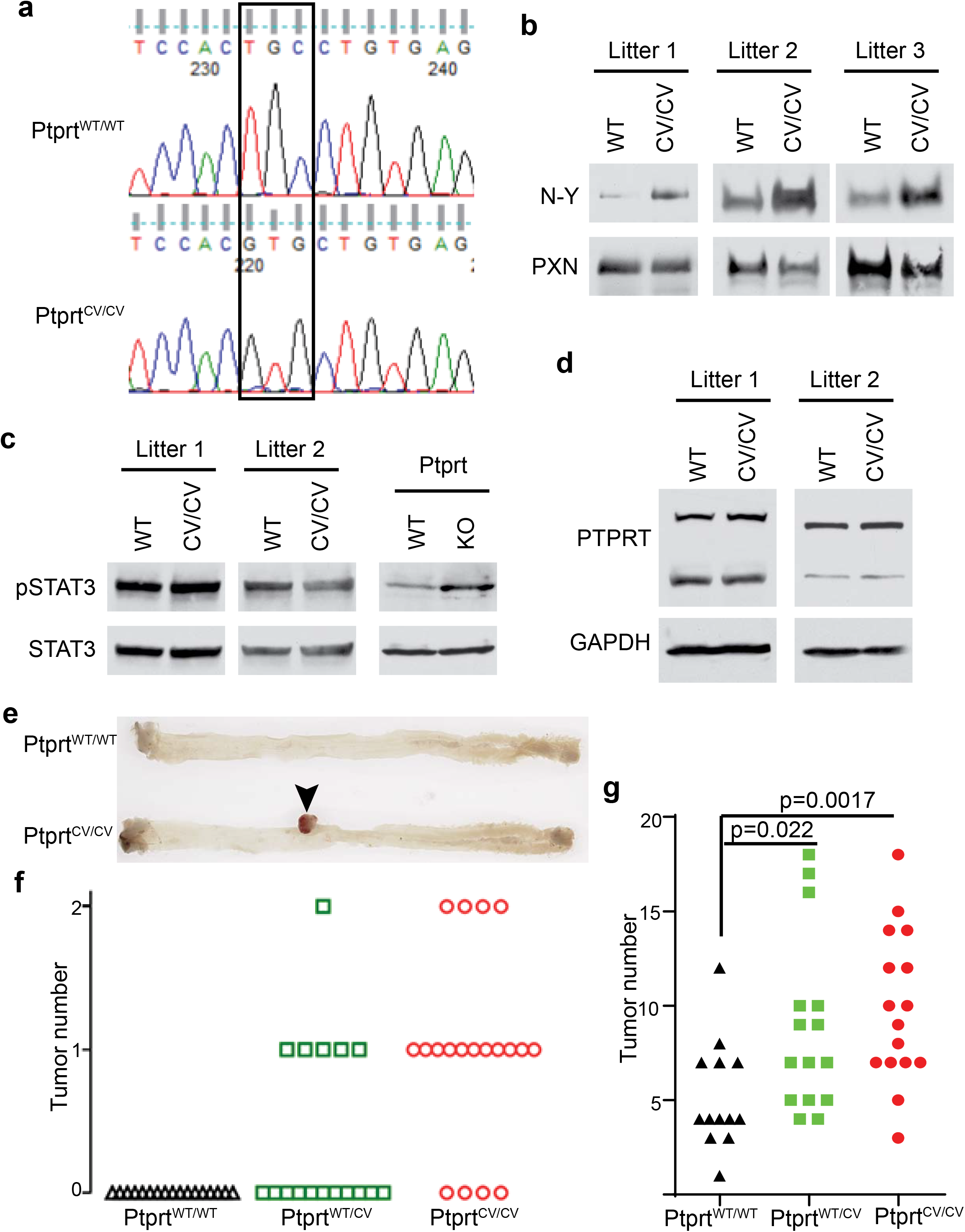
PTPRT denitrase mutant mice are susceptible to carcinogen-induced colon tumor formation. (**A**) *Generation of Ptprt^C1400V^ mutant knockin mice using CRISPR/Cas9 genome editing technology.* C57/BL6 embryos were injected with Cas9 proteins, sgRNAs and repair DNA oligonucleotides (see supplementary Fig. 3 and methods for details). Founder mice were screened for harboring the C1400V (CV) mutation. Genomic DNAs surrounding the Cas9 cut site were sequenced to ensure the presence of the C1400V mutation and absence of undesired mutations. (**B**) *Levels of paxillin tyrosine nitration are increased in Ptprt^C1400V^ mutant mice.* Paxillin proteins were immunoprecipitated from brain lysates of mice with the indicated genotype and blotted with an anti-nitrotyrosine antibody. (**C**) *The PTPRT C1400V mutation does not affect its tyrosine phosphatase activity.* Brain lysate of mice with the indicated genotypes were blotted with anti-pY STAT3 or total STAT3 antibodies. (**D**) *The PTPRT C1400V mutation does affect PTPRT protein expression levels.* Brain lysates from mice with the indicated genotypes were blotted with the indicated antibody. (**E**) & (**F**) *Ptprt^C1400V^ mutant mice are susceptible to AOM-induced colon tumor formation.* Ptprt^WT/WT^ (n=17), Ptprt^WT/C1400V^ (n=17), or Ptprt^C1400V/C1400V^ (n=19) mice were treated with six doses of AOM. Gross morphology of a tumor (indicated arrowhead) in a Ptprt^C1400V/C1400V^ mouse shown in (E) and number of tumors developed in each mouse shown in (F). (**G**) *Tumor number is increased Ptprt^C1400V^ mutant mice treated with AOM and DSS.* Tumor number in each mouse of the indicated genotype is shown. A Student’s t test was used to compare the numbers of tumor formed in Ptprt^C1400V/C1400V^ mice (n=16) or Ptprt^WT/C1400V^ (n=15) mice with number of tumor formed in Ptprt^WT/WT^ mice (n=13).

Next, we set out to determine if the denitrase activity of PTPRT contributes to its tumor suppressor function. Given that we showed previously that PTPRT knockout mice are susceptible to AOM induced colon tumor development8, we treated Ptprt^WT/WT^, Ptprt^WT/C1400V^ and Ptprt^C1400V/C1400V^ with six doses of AOM. As reported previously8, the Ptprt^WT/WT^ mice were highly resistant to AOM induced colon cancer and none developed any colon tumors (Fig. 4 E & F). In contrast, 15 of the 19 Ptprt^C1400V/C1400V^ mice developed visible colon tumors, averaging 1.2 tumors per mouse whereas 5 of 17 Ptprt^WT/C1400V^ mice developed tumors (Fig. 4F & Extended Data Fig. 4A). Moreover, in a second AOM-DSS-induced colon tumor model, Ptprt^C1400V/C1400V^ and Ptprt^WT/C1400V^ mice also developed significantly more tumors than the WT mice (Fig. 4G, Extended Data Fig. 4 B & C). Together, these data suggest that the denitrase activity of PTPRT has a tumor suppressor function.

Although protein tyrosine nitration was identified over fifty years ago^12^, the physiological relevance of this protein modification is not clear. Protein tyrosine nitration has been detected under physiological conditions as well as in diseases including cancer, atherosclerosis, myocardial infarction, chronic obstructive pulmonary disease, diabetes, Parkinson’s disease, and Alzheimer’s disease1. It is generally believed that tyrosine nitration is a byproduct of reactive oxygen/nitrogen species1. Nonetheless, several studies suggest that there exist enzymes (denitrases) that remove nitration from tyrosine residues in proteins^13-15^. The identity of those enzymes however has been elusive. We demonstrate that the second catalytic domain (D2) of PTPRT is a denitrase that removes the nitro group from tyrosine residues in paxillin and that the denitrase activity of PTPRT is associated with suppression of tumor formation in two colon cancer models. These results suggest that regulation of protein tyrosine nitration plays a critical role in tumorigenesis. Lastly, our study suggests that some, if not all, of the D2 domains of RPTPs may function as denitrases as well.

## Contributions

Z. W. conceived the idea. Y. Z., S.Z., P.Z., X. F., R.A.C., M.M. and Z.W. designed the experiments. Y. Z., S.Z., Y.H., P.Z., X. F., Y.C., and J.S. performed the experiments. Y. Z., S.Z., B.W., J.S., J.M., S.M., R.M.E., R.A.C, M.M. and Z.W. analyzed the data. Z.W., Y.Z., R.A.C., B.W., M.M., J.M., S.M, and R.M.E. wrote the manuscript.

## Acknowledgements

We thank Dr. Susann Brady-Kalnay for critical reading of the manuscript. This work was supported by NIH grants R01CA196643, R01CA127590, P50CA150964 and P30 CA043703 and Standup to Cancer Colorectal Cancer Dream Team Award (to ZW).

## Methods

### Mice

Animal experiments were approved by the Case Western Reserve University Animal Care and Use Committee. PTPRT KO mice were generated as described previously^8^. PTPRT KI mice were generated as described below. Bothe male and female mice were used.

### Generation of PTPRT^+/C1400V^ mutant mice

Fertilized C57Bl/6J mouse eggs were injected with a mixture of Cas9 mRNA, sgRNA and a 122 bp single-stranded DNA oligo. The repair oligo was designed to mutate cysteine 1400 to valine, and destroy the PAM sequence with a synonymous mutation (Extended Data Figure 3). The C1400V mutation generated a PmlI site, which was used to genotype founder animals. The knockin allele was confirmed by Sanger sequencing. Correctly targeted founder animals were used to establish and expand lines of mice by crossing them to wild type C57Bl/6J mice.

### AOM treatment

Six-week-old mice were injected intraperitoneally once weekly for 6 weeks with 10 mg/kg AOM (Sigma Chemical, St. Louis). Mice were sacrificed 24 weeks after the last AOM injection.

Colons were removed immediately, rinsed with PBS to remove fecal matter, sliced open longitudinally and analyzed for the presence of tumors. Tumors and normal colon tissue were then fixed in 10% formalin overnight to perform histological staining following paraffin embedding.

### AOM plus DSS treatment

Eight-week-old mice were injected with a single intraperitoneal 10 mg/kg body weight of azoxymethane (AOM) (Sigma-Aldrich, St Louis, MO, USA). One week later, mice were treated with 2% dextran sulfate sodium (DSS Salt Reagent Grade MW 36,000–50,000, MP Biomedicals, USA) in the drinking water for 7 days, followed by 10 days of regular water. This DSS cycle was repeated 2 times. Animals were sacrificed after 53 days. Colons were removed immediately, rinsed with PBS to remove fecal matter, sliced open longitudinally and analyzed for the presence of tumors. After that, tumors and colon tissue were fixed in 10% formalin overnight to perform histological staining after paraffin embedding.

### Cell culture and reagents

HCT116 were obtained from the American Type Culture collection (ATCC, Manassas, VA, USA). The cells were grown in McCoy’s 5A + 10% FBS and authenticated by short tandem repeat [STR] profiling. Mycoplasma contamination was routinely checked.

### Cell and tissue lysis

Cells or mouse brains were either lysed using a urea lysis buffer (10 mM Tris-HCl, 100 mM NH_2_PO_4_; 8 M urea; 5 mM Iodoacetic acid; 5mM DTT, adjust pH to 8.0) or a RIPA buffer [50 mM Tris-HCl pH 7.5; 1 mM EDTA; 150mM NaCl; 1% NP-40; 0.25% Sodium Deoxycholate; 1 mM PMSF; 1 mM Na3VO4; 20 mM NaF; protease inhibitor cocktail (Roche, Penzberg, GER); 5 mM Iodoacetic acid; 5mM DTT for co-immunoprecipitation. Western blots were performed as described previously^8^. Antibodies include anti-Nitrotyrosine (Cat No:05-233,Millipore, Temecula, CA, USA), anti-Sulfotyrosine (Cat No: 051100, Millipore, Temecula, CA, USA), anti-paxillin (Cat No: 610051, BD Biosciences, San Jose, CA, USA), anti-GAPDH (Cat No: sc-25778, Santa Cruz Biotechnology, Santa Cruz, CA, USA), anti-6X His (Cat No: ab18184, ABCAM, Cambridge, MA,USA), c-Myc (Cat No: sc-40, Santa Cruz Biotechnology, Santa Cruz, CA, USA), p-STAT3(Cat No:9145, Cell Signaling technology, Danvers, MA,USA), STAT3 (Cat No: 9139, Cell Signaling technology, Danvers, MA,USA), anti-PTPRT (Cat No:LF-MA0304, Biovendor LLC, Asheville, NC, USA) and anti-V5 (Cat No: R960-25, Invitrogen, Carlsbad,CA,USA).

### Virus infection

HCT116 cells were infected with GFP control, PTPRT WT or PTPRT ΔD2 virus for 20 hr and lysed in RIPA buffer as described above.

### Protein purification

The D1, D2 or D2CA domains of PTPRT were cloned into the pGEX vector and expressed in BL21 competent bacteria. Protein was purified using the GST-fusion purification protocol. Briefly, expression was induced for 4 hours using 0.2mM IPTG. Bacterial cells were lysed in 50 mM Tris-HCl pH 7.5; 1 mM EDTA; 150mM NaCl; 1% NP-40; 0.25% Sodium Deoxycholate; 1 mM PMSF; 1 mM Na3VO4; 20 mM NaF; 5 mM DTT and protease inhibitor cocktail, and protein was purified from lysate using Glutathione Sepharose beads and eluted with GST elution buffer (20 mM Tris-HCl pH 8.0; 1 mM EDTA; 150mM NaCl; 0.1% Triton X-100; 5 mM DTT; 20mM Glutathione; pH adjusted to 8.0). Purified protein was then dialyzed in buffer containing 50 mM Tris-HCl pH 7.5; 100mM NaCl; 10 % Glycerol; 5 mM DTT using a slide-a-lyzer dialysis cassette (Thermo Fisher Scientific, Waltham, MA, USA).

The different fragments of paxillin were cloned into the pET28a vector and expressed in BL21 competent bacteria. Protein was purified using the Qiaexpressionist (Qiagen, Valencia, CA, USA) protocol. Briefly, expression was induced overnight using 0.1mM IPTG, cells were lysed in 50 mM NH_2_PO_4_, 300 mM NaCl, 10 mM imidazole, pH 8.0. Protein was purified from the lysate using Ni-NTA agarose. Purified protein was then dialyzed in PBS using a slide-a-lyzer dialysis cassette (Thermo Fisher Scientific, Waltham, MA, USA).

### Nitration of substrate

Recombinant paxillin protein (200 ng/μl) was treated with 2 mM 5,5’-Dithiobis(2-nitrobenzoic acid) and 0.1 mM Diethylenetriaminepentaacetic acid (DTPA) in pH 8.0, 150 mM Tris-HCl buffer and incubated for 30 min at 25°C to protect the sulfhydryl groups of cysteine residues. Excess reagent and reaction byproducts were removed and the buffer was exchanged to 50 mM Tris-HCl/0.1mM DTPA, pH8.0 using a Microcon-10kDa centrifugal filter unit (Millipore). The protein was then nitrated by adding tetranitromethane (TNM) to a final concentration of 1 mM and incubating for 10 min at 25°C. The nitrated *S*-protected protein was treated with 10 mM DTT for 1 h at 25°C to de-protect the sulfhydryl groups of cysteine residues and then dialyzed against 50 mM Tris-HCl (pH 7.5) containing 100 mM NaCl at 4°C overnight.

### In vitro denitrase assay

PTPRT recombinant proteins (200 ng) were mixed in buffer containing 25mM Tris-HCl, 50mM NaCl and 1:7 diluted HCT116 cell lysate filtered through a 3 KD cut-off spin column to get rid of large molecular weight materials, which provides necessary co-factors. The reactions were pre-incubated for 30 min at 25°C. Nitrated-paxillin substrates (200 ng) were then added in a total reaction volume of 200 μl and incubated for 15 minutes to 2 hr at 30°C. The mixtures were resolved 10% SDS-PAGE, and proteins were either transferred to nitrocellulose membranes for Western blot analyses or excised for mass spectrometry analyses as described below.

### Enzyme kinetics assay

Various concentrations (2.688 mM, 1.344 mM, 0.672 mM, 0.336 mM, 0.168 mM, or 0 nM) of synthesized paxillin N-Y404 peptide [RDGQP (3-NO2-Tyr) CEKDY; Genscript, Piscataway, NJ] were incubated with 41 nM of WT or mutant PTPRT recombinant proteins to perform in vitro denitrase assay as described above. Amounts of N-Y paxillin peptide were measured by absorbance of nitrotyrosine at a wavelength of 360 using a SpectraMax i3x Multi-Mode microplate reader (Molecular Devices, Sunnyvale, CA, USA). Amounts of paxillin N-Y404 peptide were calculated according a standard linear absorbance curve [Peptide concentration (mM) = (A360 - 0.00295)/1.434]. The rate of denitration of N-Y404 paxillin peptide was determined according to the decrease in A360nm in 30 min. It was established in separate experiments that the 30-min time point was within the linear time course of the reaction, and that 41 nM PTPRT was within the range of linear dependence of the rate on enzyme concentration. Enzymatic kinetics were calculated using the Prism Software (Graphpad).

### In-gel digestion

SDS-PAGE gel bands containing paxillin protein were subjected to in-gel digestion with sequencing grade modified trypsin (Promega, WI). Briefly, gel pieces excised from a SDS-PAGE were first washed with 50% acetonitrile in 50 mM ammonium bicarbonate, and then dehydrated with acetonitrile. Before an overnight proteolytic digestion, proteins were reduced with 100 mM DTT in 100 mM ammonium bicarbonate at 25 °C for 30min, and alkylated with 250 mM iodoacetamide in 100 mM ammonium bicarbonate, and incubate for 30 min at 25°C. After proteolytic digestion, peptides were extracted from the gel with 50% acetonitrile and then re-suspended in 0.1% formic acid after being dried completely under the Speed Vacuum.

### Liquid chromatography–tandem mass spectrometry

Liquid chromatography–tandem mass spectrometry (LC-MS/MS) was carried out using a Waters nanoAcquity UPLC system (Waters, Taunton, MA, USA) interfaced to Orbitrap Elite Hybrid Mass Spectrometer (Thermo Electron, San Jose, CA, USA). Peptides were chromatographed on a reversed-phase 0.075 × 250-mm C18 Waters nanoACQUITY UPLC analytical column (Waters) using a linear gradient of acetonitrile from 2% to 35% over 60 min followed by from 35% to 90% in 15 min in aqueous 0.1% formic acid at a flow rate of 300 nL/min. The eluent was directly introduced into the mass spectrometer operated in a data-dependent MS to MS/MS switching mode, with the 20 most intense ions in each MS scan subjected to MS/MS analysis. The full MS scan was performed at a resolution of 120,000 (full width at half-maximum) in the Orbitrap detector, and the MS/MS scans were performed in the ion trap detector in collision-induced dissociation mode with a normalized collision energy of 35 %. The obtained data were submitted for search against customized paxillin protein database using a Mascot Daemon database search engine. Carbamidomethylation of Cys, oxidation of Met and nitration of Tyr were selected as variable modifications. The mass tolerance was set as 10 ppm for precursor ions and 0.8 Da for product ions. The candidates of nitrated peptides and corresponding denitrated peptides suggested by the search engine were further verified by manual interpretation of the MS/MS spectra.

### Statistical analysis

We applied the Student’s *t* test to compare the means between two groups, assuming unequal variances.

### Data availability

The data that support the findings of this study are available from the corresponding author upon request.

## Extended data

**Extended Data Figure 1.**
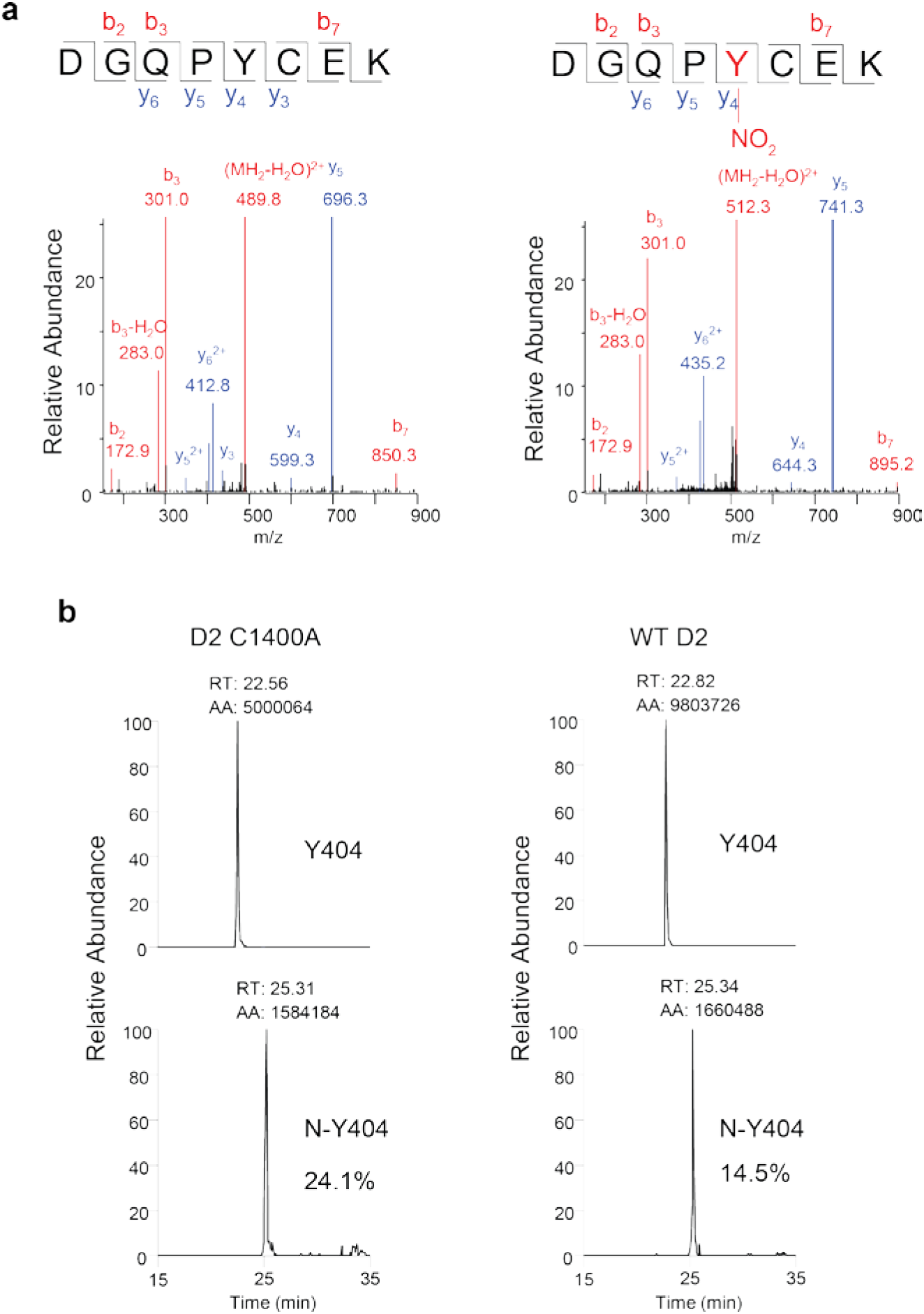
Mass Spectrometry analysis of a nitro-Y404 paxillin peptide. (**A**) MS/MS spectra of a nitro-Y404 paxillin peptide (DGQPYCEK, residue 400-407) and the same peptide denitrated by WT PTPRT D2 recombinant protein are shown. Doubly charged precursor ions from the nitrated (m/z 521.2) and denitrated (m/z 498.7) peptides were subjected to collision-induced dissociation. Cys405 in the peptide was carbamidemethylated. A series of b and y series fragment ions that verify the chemical structures of the peptides were observed. (B) Quantification of the Y404-nitrated and denitrated peptide when M-2 paxillin fragments were incubated with either WT or C1400A PTPRT D2 recombinant protein. The ions for the nitrated (N-Y404) and denitrated (Y404) peptides were extracted from the total ion chromatogram for precursor ions, and the chromatographic peak areas between nitrated and denitrated peptide ions were compared to calculate the fraction of nitrated peptide. The results indicated that the extent of nitration at Y404 was decreased after incubating with WT PTPRT D2.

**Extended Data Figure 2.**
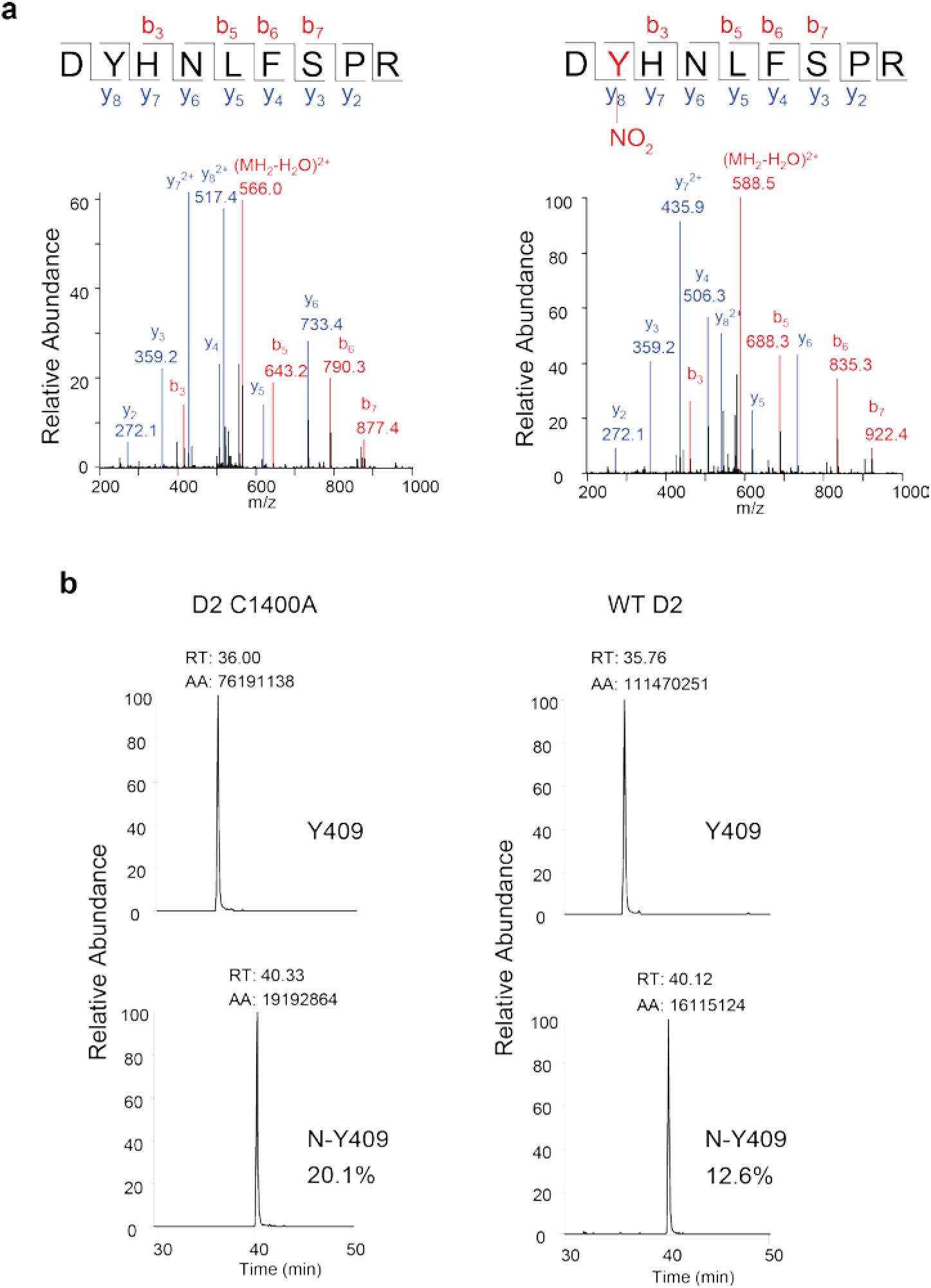
Mass Spectrometry analysis of a nitro-Y409 paxillin peptide. (**A**) MS/MS spectra of a nitro-Y409 paxillin peptide (DYHNLFSPR, residue 408-416) and the same peptide denitrated by WT PTPRT D2 recombinant protein are shown. Doubly charged precursor ions from the nitrated (m/z 597.2) and denitrated (m/z 574.7) peptides were subjected to collision-induced dissociation. A series of b and y series fragment ions that verify the chemical structures of the peptides were observed. (B) Quantification of the Y409-nitrated and denitrated peptides when the M-2 paxillin fragments were incubated with either WT or C1400A PTPRT D2 recombinant protein. The ions for the nitrated (N-Y409) and denitrated (Y409) peptides were extracted from the total ion chromatogram for precursor ions, and the chromatographic peak areas between nitrated and denitrated peptide ions were compared to calculate the fraction of nitrated peptide. The results indicated that the extent of nitration at Y404 was decreased after incubating with WT PTPRT D2.

**Extended Data Figure 3.**
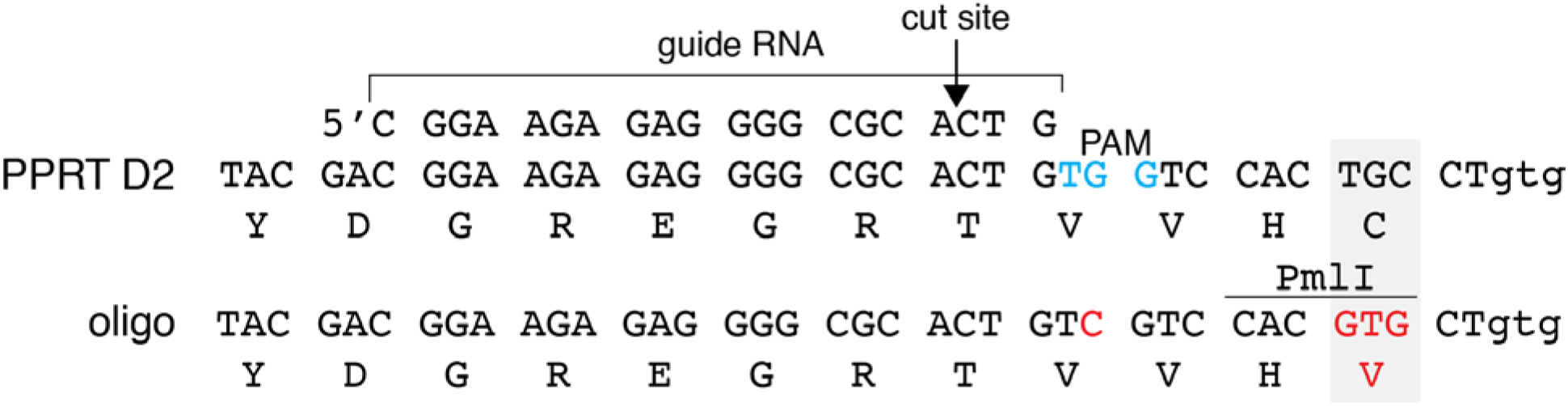
Strategy to generate a C1400V knockin mutation in mice with CRISPR/Cas9. The conserved cysteine residue and the knockin mutation to valine are highlighted in grey. The targeting oligo was designed to create a PmlI restriction site for genotyping mice and a synonymous mutation to destroy the PAM sequence. Arrow indicates the Cas 9 cutting site.

**Extended Data Figure 4.**
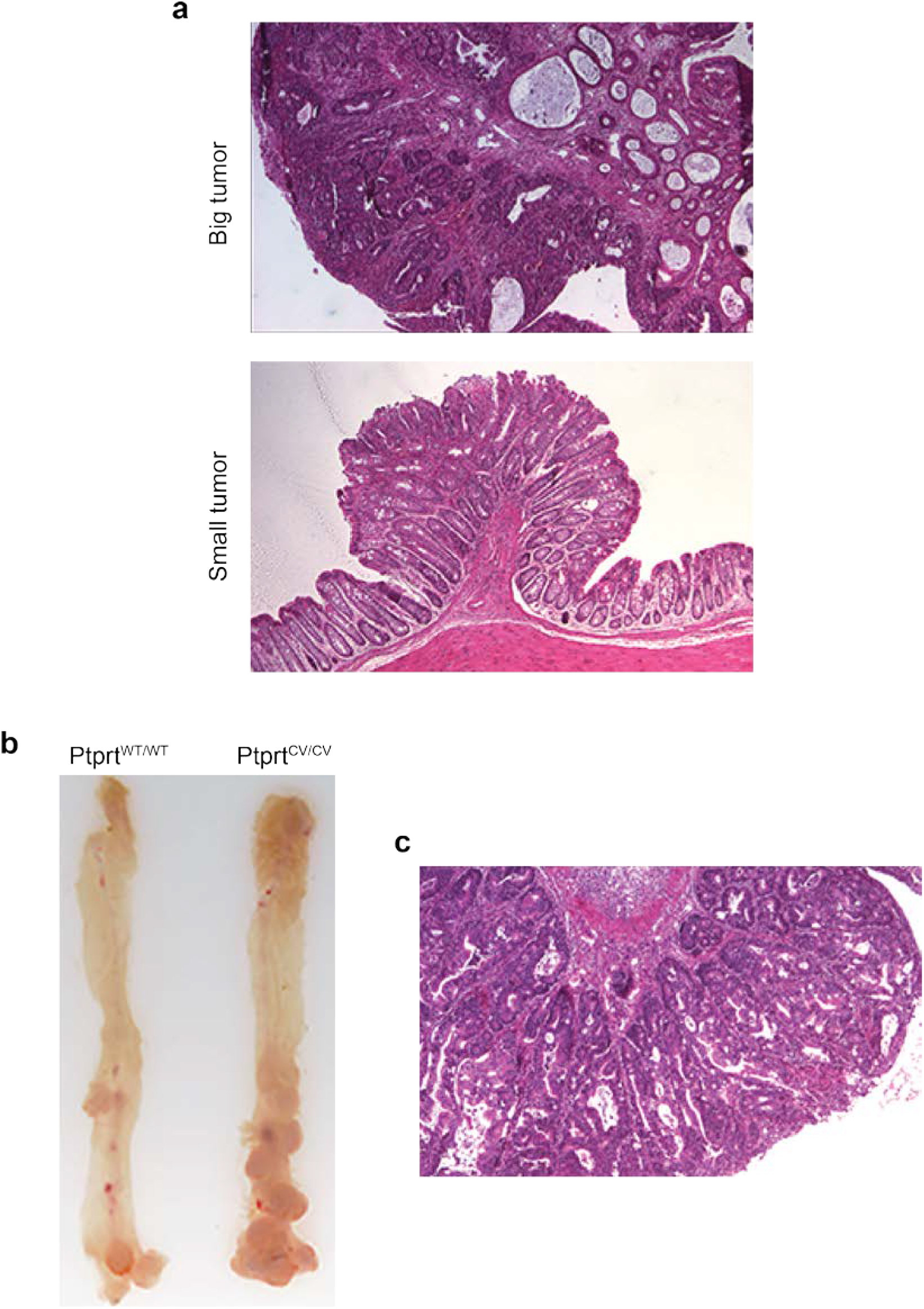
Tumors developed in Prprt^C1400V/C1400C^ mice. (**A**) H&E staining of representative tumors developed in the AOM-induced colon tumor model. (**B**) & (**C**) Tumors developed in the AOM-DSS induced tumor model. Photography of colon of mice with the indicated genotype shown in (B) and H&E staining of a tumor shown in (C).

## References

1 Radi, R. Nitric oxide, oxidants, and protein tyrosine nitration. Proceedings of the National Academy of Sciences of the United States of America 101, 4003–4008, doi:10.1073/pnas.0307446101 (2004).

2 Wang, Z. et al. Mutational analysis of the tyrosine phosphatome in colorectal cancers. Science 304, 1164–1166 (2004).

3 Zhao, S., Sedwick, D. & Wang, Z. Genetic alterations of protein tyrosine phosphatases in human cancers. Oncogene 34, 3885–3894, doi:10.1038/onc.2014.326 (2015).

4 Alonso, A. et al. Protein tyrosine phosphatases in the human genome. Cell 117, 699–711 (2004).

5 Chen, M. J., Dixon, J. E. & Manning, G. Genomics and evolution of protein phosphatases. Science signaling 10, doi:10.1126/scisignal.aag1796 (2017).

6 Frankson, R. et al. Therapeutic Targeting of Oncogenic Tyrosine Phosphatases. Cancer research 77, 5701–5705, doi:10.1158/0008-5472.CAN-17-1510 (2017).

7 Tonks, N. K. & Neel, B. G. Combinatorial control of the specificity of protein tyrosine phosphatases. Current opinion in cell biology 13, 182–195 (2001).

8 Zhao, Y. et al. Identification and functional characterization of paxillin as a target of protein tyrosine phosphatase receptor T. Proceedings of the National Academy of Sciences of the United States of America 107, 2592–2597, doi:0914884107 [pii]10.1073/pnas.0914884107 (2010).

9 Zhao, Y. et al. Regulation of paxillin-p130-PI3K-AKT signaling axis by Src and PTPRT impacts colon tumorigenesis. Oncotarget, doi:10.18632/oncotarget.10654 (2016).

10 Reinders, J. & Sickmann, A. Modificomics: posttranslational modifications beyond protein phosphorylation and glycosylation. Biomolecular engineering 24, 169–177, doi:10.1016/j.bioeng.2007.03.002 (2007).

11 Zhang, X. et al. Identification of STAT3 as a substrate of receptor protein tyrosine phosphatase T. PNAS 104, 4060–4064, doi:10.1073/pnas.0611665104 (2007).

12 Sokolovsky, M., Riordan, J. F. & Vallee, B. L. Tetranitromethane. A reagent for the nitration of tyrosyl residues in proteins. Biochemistry 5, 3582–3589 (1966).

13 Kamisaki, Y. et al. An activity in rat tissues that modifies nitrotyrosine-containing proteins. Proceedings of the National Academy of Sciences of the United States of America 95, 11584–11589 (1998).

14 Deeb, R. S. et al. Characterization of a cellular denitrase activity that reverses nitration of cyclooxygenase. American journal of physiology. Heart and circulatory physiology 305, H687–698, doi:10.1152/ajpheart.00876.2012 (2013).

15 Irie, Y., Saeki, M., Kamisaki, Y., Martin, E. & Murad, F. Histone H1.2 is a substrate for denitrase, an activity that reduces nitrotyrosine immunoreactivity in proteins. Proceedings of the National Academy of Sciences of the United States of America 100, 5634–5639, doi:10.1073/pnas.1131756100 (2003).

